# Chromosome assembly of large and complex genomes using multiple references

**DOI:** 10.1101/088435

**Authors:** Mikhail Kolmogorov, Joel Armstrong, Brian J. Raney, Ian Streeter, Matthew Dunn, Fengtang Yang, Duncan Odom, Paul Flicek, Thomas Keane, David Thybert, Benedict Paten, Son Pham

## Abstract

Despite the rapid development of sequencing technologies, assembly of mammalian-scale genomes into complete chromosomes remains one of the most challenging problems in bioinformatics. To help address this difficulty, we developed Ragout, a reference-assisted assembly tool that now works for large and complex genomes. Taking one or more target assemblies (generated from an NGS assembler) and one or multiple related reference genomes, Ragout infers the evolutionary relationships between the genomes and builds the final assemblies using a genome rearrangement approach. Using Ragout, we transformed NGS assemblies of 15 different *Mus musculus* and one *Mus spretus* genomes into sets of complete chromosomes, leaving less than 5% of sequence unlocalized per set. Various benchmarks, including PCR testing and realigning of long PacBio reads, suggest only a small number of structural errors in the final assemblies, comparable with direct assembly approaches. Additionally, we applied Ragout to *Mus caroli* and *Mus pahari* genomes, which exhibit karyotype-scale variations compared to other genomes from the *Muridae* family. Chromosome color maps confirmed most large-scale rearrangements that Ragout detected.

## Introduction

The year 2001 marked an important step in genome biology with the release of the first near complete human genome [Lander et al., 2001, Venter et al., 2001]. Since then, numerous near complete mammalian genome sequences have been made available [Pontius et al., 2007, Church et al., 2009, Scally et al., 2012]. These finished genomes, while being expensive to produce, have greatly advanced the field of comparative genomics and provided many new insights to our understanding of mammalian evolution. The initial achievement was quickly followed by the era of high throughput sequencing technologies – next generation sequencing (NGS). These cost-effective technologies are allowing many sequencing consortia to explore genomes from a large number of species [Jarvis et al. 2014].

Recently, new de novo assembly algorithms have been developed to combine high-throughput short read sequencing data with long single molecule sequencing reads or jumping libraries to completely assemble bacterial genomes (one chromosome into one contig) [Koren and Phillippy, 2015]. By contrast, complete assembly of mammalian genomes using current short-read sequencing technologies remains a formidable problem, since such genomes are larger and have more complicated repeat structures. Most current mammalian assemblies produced by NGS-assemblers [Butler et al., 2008, Simpson et al., 2009] contain thousands to hundreds of thousands of contigs/scaffolds and provide limited value for comparative genomics as constructed syntenic regions are highly fragmented. Recently, some studies [Gordon et al., 2016, Chaisson et al., 2015] have applied long read technologies (Pacific Biosciences, 10x Genomics, Dovetail) to improve the assembly of larger genomes. However, the cost of generating high throughput long reads is still much higher than the cost of generating short read libraries.

Since many complete genomes are now available, an alternative approach is to use these genomes to guide the assembly of the target (assembled) genome, in a method called ‘reference-assisted assembly’ [Gnerre et al., 2009]. In such methods, the information from a closely-related reference genome is used by an NGS-assembler for resolving complicated genomic structures, such as repeats or low-coverage regions. This technique was implemented in a number of assemblers/scaffolders [Zerbino and Birney, 2008, Peng et al., 2010, Gnerre et al., 2011, Iqbal et al., 2012] and proved to be valuable when a close reference is available. Another common approach is to align pre-assembled contigs of the target genome against the reference, and order them according to their positions in the reference genome [Richter et al., 2007, Rissman et al., 2009]. However, for both approaches, simplistic modeling, in which each breakpoint is treated independently, still introduces many misassembly errors when structural variations between the reference and target genomes are present.

To improve over the single reference genome approach Kim et al. [2013] introduced the RACA tool, which made an important step toward reliable reconstruction of the target genome by analyzing the structure of multiple outgroup genomes in addition to a single reference. This approach proved to be valuable, since consistent adjacency information across multiple outgroups proved a more powerful predictor of adjacencies in the target assembly than single genomes alone. Similar to other genome rearrangement approaches, RACA relies on the decomposition of the input sequences into a set of synteny blocks – long and conservative genomic regions with respect to micro-rearrangements. However, given the lack of synteny block reconstruction tools for mammalian sequences (only tools for pairwise comparisons are available), RACA reconstructs synteny blocks by aligning all input sequences against a single reference genome. This approach is biased towards the reference genome and in some cases, cannot detect synteny blocks [Pham and Pevzner, 2010]. As a result, RACA, while showing its improvements over other tools that use a single reference, still generates misassemblies and gaps in its final scaffolds.

This work presents Ragout, an algorithm (and a software package) that attempts to address all of the above challenges in reference-assisted assembly for mammalian genomes. Ragout combines Progressive Cactus, a multiple whole genome aligner [Paten et al., 2011], with a new graph simplification algorithm to decompose the input sequences into multi-scale synteny blocks and an iterative algorithm for finding the missing adjacencies. Ragout utilizes multi-scale synteny blocks to separate large structural variations from smaller polymorphisms, and it also minimizes the number of gaps and mis-joins in the final scaffolds. Additionally, a 2-break rearrangement model [Alekseyev and Pevzner, 2009] is used to distinguish between target-specific rearrangements and chimeric adjacencies.

## Results

### Synteny blocks

Nucleotide-level alignments between diverged genomes contain millions of small variations. To analyze karyotype-level rearrangements, studies typically use lower-resolution alignments. Such alignments are described as a set of coarse synteny blocks [Kent et al., 2003, Pevzner et al., 2003], each such synteny block being a set of strand-oriented chromosome intervals in the set of genomes being compared that represents the homology relationship between large segments of the genomes. In this study, we also use synteny blocks to separate large structural variations from small polymorphisms. However, we take a hierarchical approach, with multiple sets of synteny blocks, each defined at a different resolution, from the coarsest, karyotype level all the way down to the fine-grained base level. To create the hierarchy, we use the principles developed by the Sibelia tool [Minkin et al., 2013], which can create such a hierarchy for bacterial genomes, but adapted with a graph simplification algorithm for constructing synteny blocks from a multiple genome alignment file in HAL format [Hickey et al., 2013], produced by Progressive Cactus [Paten et al., 2011]. The algorithm starts from synteny blocks of the highest resolution (local sequence alignments) and then iteratively merges them into larger blocks if they are structurally concordant. Thus, each coarse synteny block has multiple predecessors, which defines the hierarchy (see Methods section for the details).

### A rearrangement approach for genome assembly

At each level of resolution, the Ragout algorithm decomposes the input genomes into a set of signed strings of synteny blocks, such that when the set of strings is concatenated together it forms a signed permutation of blocks. While each contiguous reference chromosome is transformed into a single sequence of signed synteny blocks, each chromosome in the target assembly corresponds to multiple sequences of synteny blocks because of the assembly fragmentation. As a result, some information about the adjacencies between synteny blocks in the target genome is missing. Ragout infers a phylogenetic relationship between the genomes and constructs a breakpoint graph from the sets of synteny block strings, and then uses a rearrangement approach to infer these missing adjacencies. Additionally, a 2-break rearrangement model [Alekseyev and Pevzner, 2009] is used to identify chimeric adjacencies in the input contigs/scaffolds and distinguish them from real rearrangements. All these steps are performed iteratively, starting from the lowest to the highest synteny block resolution. In each iteration, scaffolds in the previous step are merged with scaffolds in the current step to refine the resulting assembly. Ragout further applies a “context-matching” algorithm to resolve repeats and put them back into the final scaffolds (see Figure 1 for an algorithm overview and the Methods section for details).

**Figure 1.**
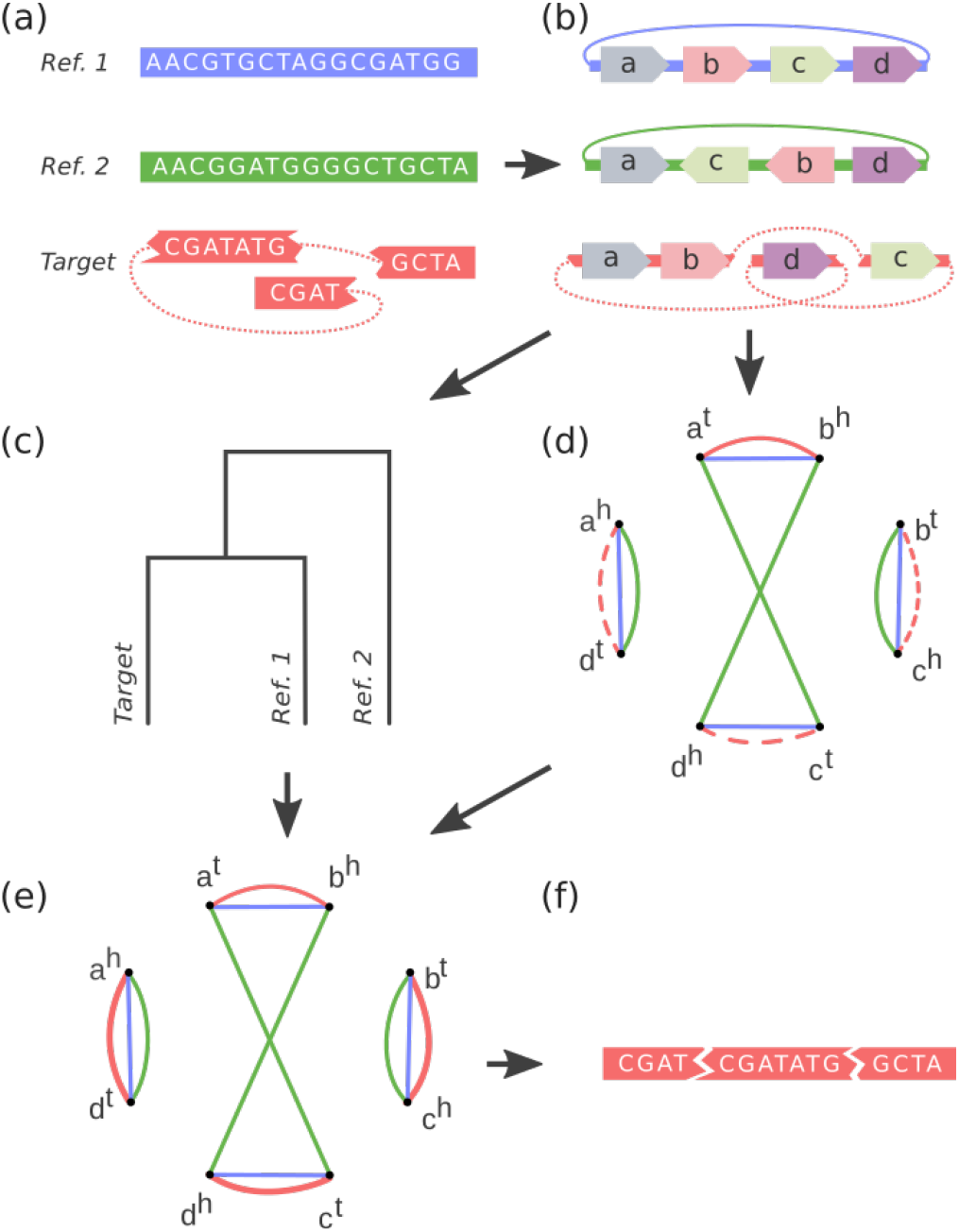
An overview of the Ragout algorithm. Reference and target nucleotide sequences (a) are first decomposed into the permutation of synteny blocks. Using the synteny information (b) and the phylogenetic relationships between the input genomes (c), it constructs an incomplete breakpoint graph (d), in which each synteny block *b* corresponds to two vertices, representing its head and tail (b^h^ and b^t^). The graph represents adjacency information between synteny blocks. Afterwards, the algorithm recovers missing adjacencies in the breakpoint graph (e) and uses them to merge corresponding contigs/scaffolds into chromosomes (f).

### Benchmarking Ragout and RACA on simulated datasets

We simulated multiple datasets with extensive structural rearrangements to benchmark the Ragout algorithm and compared its performance against RACA. We took human chromosome 14 (GRCh37/hg19 version) as an ancestral genome and chose a set of breakpoints such that they divide the chromosome into intervals of exponentially distributed length [Pevzner et al., 2003], in approximate concordance with empirical data. Then, we modeled structural rearrangements (inversion, translocation and gene conversion) with breakpoints randomly drawn from the defined breakpoint set. These rearrangements were uniformly distributed on the branches of the phylogenetic tree. Next, we modeled the NGS assembly process by fragmenting the target genome. Since each breakpoint in mammalian genomes is close to repetitive elements [Pevzner et al., 2003, Brueckner et al., 2012], we marked half of the repeats that are located near the breakpoints as “unresolved” by the assembler and fragment the genome in the corresponding positions. We further applied additional fragmentation at random positions to make the dataset more complicated. Finally, to mimic chimeric scaffolds generated in typical NGS studies [Kim et. al., 2013], we randomly joined together 5% of the target fragments. Using this setting, we simulated three reference genomes and a target genome consisting of approximately 5000 fragments with a mean length of 18,000 bp. The corresponding phylogenetic tree is shown in Figure 2(a). As RACA requires paired-end reads as input, for each target genome we sampled 30 million reads of length 70 bp using the wgsim software [Li, 2013].

We then benchmarked Ragout and RACA on datasets simulated as described above with different number of rearrangements (ranging from 50 to 500, which also corresponded to the breakpoint reuse rates of 5% and 50%, respectivelty). To benchmark the chimera detection module, we performed extra Ragout runs with the following modifications. The first extra run is called *permissive* with the chimera detection module turned off. The second extra run is denoted as *conservative*, in which all unsupported target adjacencies are broken (to mimic the common mapping approach in reference-guided assembly).

**Figure 2.**
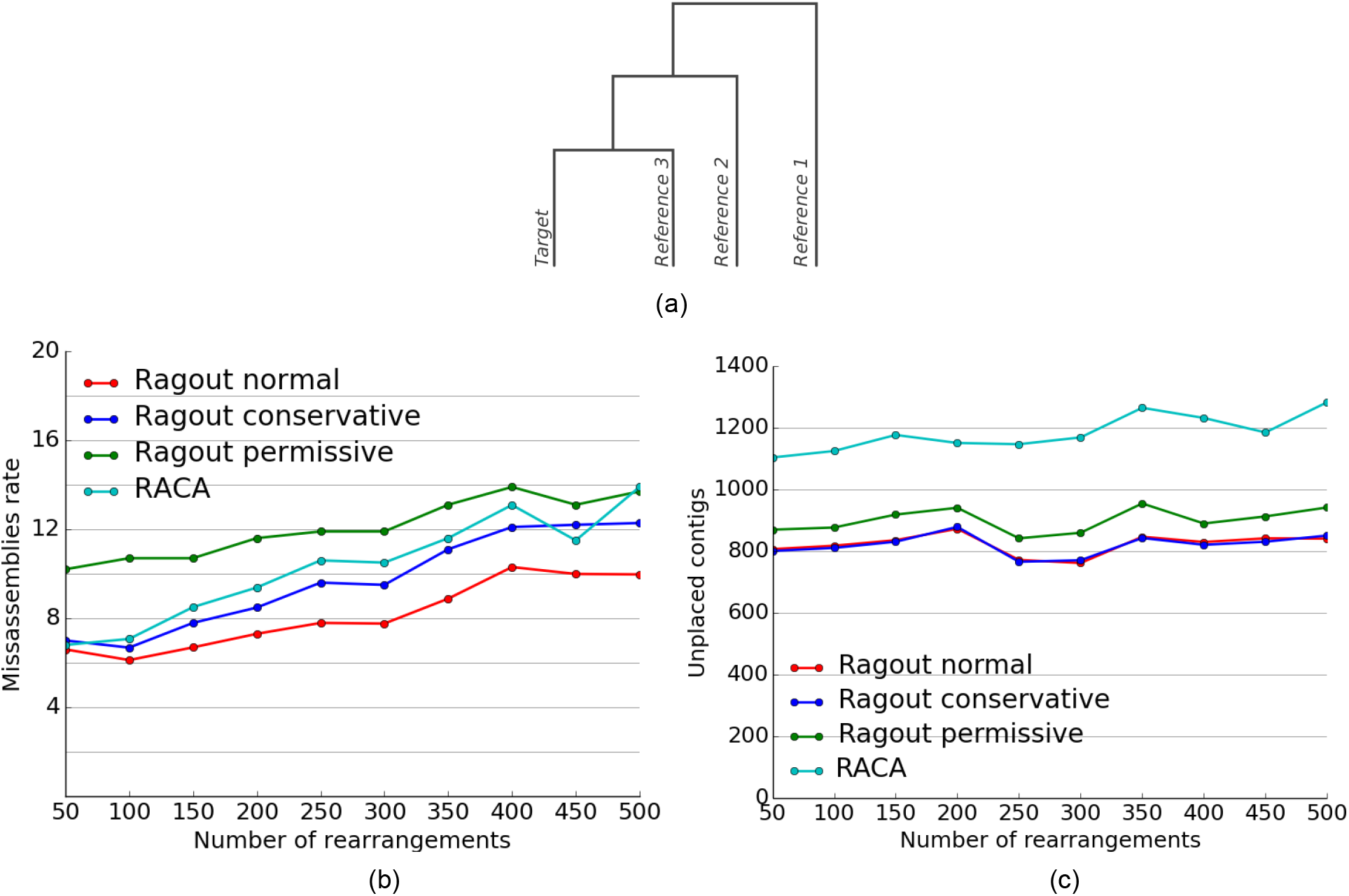
(a) The phylogenetic tree structure for the simulated datasets. All tree branches have equal length. Misassembly rates (b) and the number of unplaced contigs (c) depend on the number of rearrangements. Different algorithms are compared using the complete set of three references.

Given a resulting set of chromosomes, we call an adjacency (a pair of consecutive contigs) *correct* if these contigs have the same sign and their original positions in the target genome are adjacent (allowing jumps through unplaced fragments which were not included into the final scaffolds). Otherwise, the adjacency is called *erroneous*. For each run, we measured the error rate as the number of erroneous adjacencies divided by the total number of adjacencies. The computed error rates as well as the statistics of unplaced contigs are shown on the Figure 2. As was expected, the normal Ragout strategy, which keeps a subset of target-specific adjacencies (rather than all/none of them), produces fewer errors compared to the naive strategies. Interestingly, RACA performance was bounded by the conservative and permissive Ragout strategies for the most of the datasets. The normal and conservative strategies resulted in almost the same number of unplaced contigs. However, this number is actually higher for the permissive strategy, which is a consequence of including chimeric adjacencies in the final scaffolds. RACA produced a higher number of unplaced contigs compared to all Ragout strategies. Detailed assembly statistics for the first dataset with 50 rearrangements are given in Supplementary Table 4. We additionally benchmarked Ragout on incomplete sets of references, for which the results are given in Supplementary Materials S5. Also see Supplementary Materials S2 for comparison of Ragout and RACA using the comprehensive dataset of multiple human genome assemblies [Salzberg et al., 2012].

### Assembly of 15 *Mus musculus* and one *Mus spretus* genomes

We applied Ragout to 15 *Mus musculus* and one *Mus spretus* genomes, which included 12 laboratory strains (*129S1/SvimJ, A/J, AKR/J, BALB/cJ, C3H/HeJ, C57BL/6NJ, CBA/J, DBA/2J, FVB/NJ, LP/J, NOD/ShiLtJ, NZO/H1LtJ*) and four wild-derived strains (*WSB/EiJ, PWK/PhJ, SPRET/EiJ, CAST/EiJ*) [Keane et al., in preparation]. The initial NGS assembly was performed using SGA [Simpson et al., 2009]. Scaffolding was performed with SOAPdenovo2 [Luo et al., 2012] using multiple paired-read and mate-pair libraries with insert sizes 3, 6 and 10 kbp, 40 kbp fosmid ends for eight strains, and BAC ends for NOD/ShiLtJ. Additionally, for the three most divergent genomes (*PWK/PhJ, SPRET/EiJ and CAST/EiJ)* scaffolding based on a Dovetail protocol [Putnam et al., 2016] was applied. We used the *C57BL/6J M. musculus* strain as a single reference, as all target genomes have the same karyotype and show a good structural similarity with the *C57BL/6J* reference: On average 254 adjacent synteny block pairs longer than 10 kbp from the target NGS assemblies were not adjacent in the *C57BL/6J* reference (which correspond to large putative rearrangements). For each target strain, Ragout produced a complete set of chromosomes with the expected large-scale structure. Some assemblies also included short unlocalized fragments (homologous to the corresponding sequences in *C57BL/6J*) or MT chromosomes. The statistics of assembly results are given in Table 1. The unplaced sequence for each assembly comprised less than 5% of the total length. We also estimated the number of missing exons as less than 2% for each assembly (see below). On average, 49 adjacencies between synteny blocks larger than 10 kbp from the assembled chromosomes were not present in the *C57BL/6J* reference (excluding the three most divergent genomes).

We used multiple sets of PacBio reads that were available for the three most divergent genomes (*PWK/PhJ, SPRET/EiJ and CAST/EiJ*) to estimate the structural accuracy of the assembled chromosomes. The samples included whole genome sequence data at approximately 0.5-1x coverage and mean read length of approximately 3,000 bp as well as RNA sequencing data from liver and spleen for each of the three strains. We classified each adjacency (a pair of consecutive NGS fragments on a Ragout chromosome) as *covered* if both fragments have read alignments of at least 500 bp, or *uncovered* otherwise (47%, 43% and 55% adjacencies were covered for the *PWK/PhJ, SPRET/EiJ and CAST/EiJ* genomes, respectively). We call an adjacency *validated* if there is at least one read that has alignments longer than 500 bp on both parts of the adjacency with a correct orientation. We then calculated the validated adjacency ratio as the number of validated adjacencies divided by the number of covered adjacencies (Figure 3 shows the validated adjacency ratio as a function of the maximum gap size of an adjacency). As expected, longer gaps were harder to validate as the chance of being covered with a single read decreases. Interestingly, genomes that were closer to the *C57BL/6J* reference had more validated adjacencies, which is explained by the increasing structural divergence between the reference and the target assembly. Additionally, since the exome data had lower coverage, fewer adjacencies could have been validated using that dataset. The probability of a correct and covered adjacency without a gap not being validated by the reads of length 3,000 kbp at 1x coverage could be estimated as 15%, which was in agreement with the experimantal data (see Supplementary Material S7 for the details).

**Table 1.** Summary 16 mouse strain assemblies. Lost exons are defined as protein-coding exons which do not align on chromosomes or have a better alignment on unplaced sequence than on chromosomes. The total number of protein-coding exons in the database was 356,151. The fraction of unplaced sequence does not exceed 5% for each genome. Similarly, the fraction of lost exons does not exceed 2%. The number of broken (specific) adjacencies corresponds to the adjacencies between synteny blocks longer than 10 kbp in the assembled chromosomes that were not observed in the *C57BL/6J* reference and classified as chimeric and removed (true rearrangements and included into the final chromosomes).

| Strain | Scaffolds | Scaffolds length (mb) | Unplaced sequence (mb) | Lost exons | Broken adjacencies | Specific adjacencies |
| --- | --- | --- | --- | --- | --- | --- |
| 129S1/SvimJ | 21 | 2,694 | 119 | 6,035 | 145 | 26 |
| A/J | 22 | 2,590 | 103 | 6,283 | 212 | 52 |
| AKR/J | 22 | 2,633 | 109 | 6,318 | 124 | 32 |
| BALB/cJ | 22 | 2,594 | 92 | 5,108 | 228 | 75 |
| C3H/HeJ | 22 | 2,666 | 107 | 5,328 | 71 | 33 |
| C57BL/6NJ | 23 | 2,765 | 108 | 4,737 | 275 | 65 |
| CAST/EiJ | 22 | 2,868 | 115 | 6,203 | 74 | 249 |
| CBA/J | 21 | 2,872 | 115 | 5,440 | 189 | 58 |
| DBA/2J | 21 | 2,576 | 105 | 5,682 | 170 | 39 |
| FVB/NJ | 21 | 2,560 | 112 | 7,040 | 88 | 38 |
| LP/J | 21 | 2,695 | 123 | 5,384 | 125 | 53 |
| NOD/ShiLtJ | 21 | 2,922 | 109 | 5,680 | 125 | 47 |
| NZO/H1LtJ | 23 | 2,655 | 108 | 5,320 | 69 | 31 |
| PWK/PhJ | 21 | 2,588 | 108 | 5,487 | 87 | 347 |
| SPRET/EiJ | 21 | 2,652 | 116 | 5,558 | 116 | 659 |
| WSB/EiJ | 22 | 2,662 | 145 | 6,610 | 57 | 41 |

Finally, we benchmarked the ability of Ragout to preserve target-specific rearrangements that are not observed in reference genomes. We used 688 PCR primer pairs available from the previous studies [Yalcin et al., 2012] that surround structural variations in different *M. musculus* genomes [Keane et al., 2014]. For each genome we extracted primer pairs that align on a single NGS scaffold with a variation in distance with respect to the *C57BL/6J* reference (on average, 496 primer pairs were chosen). For each such pair we compared the alignment distances between the NGS assembly and the Ragout chromosomes. Surprisingly, while Ragout corrected many structural misassemblies (see Table 1), all these target-specific structural variations were preserved.

*We also compared Ragout performance against RACA using three Mus genomes for which long PacBio reads were available (PWK/PhJ, SPRET/EiJ and CAST/EiJ)*. Similarly to Ragout runs, RACA runs were performed using one reference (*C57BL/6J*) as well as all available paired-end and mate-pair libraries. Ragout chromosomes consistently showed less split transcripts and transcripts with wrong orientation, while the number PacBio reads with correct alignment orientations was higher (see Supplementary Material S1 for the detailed comparison).

**Figure 3.**
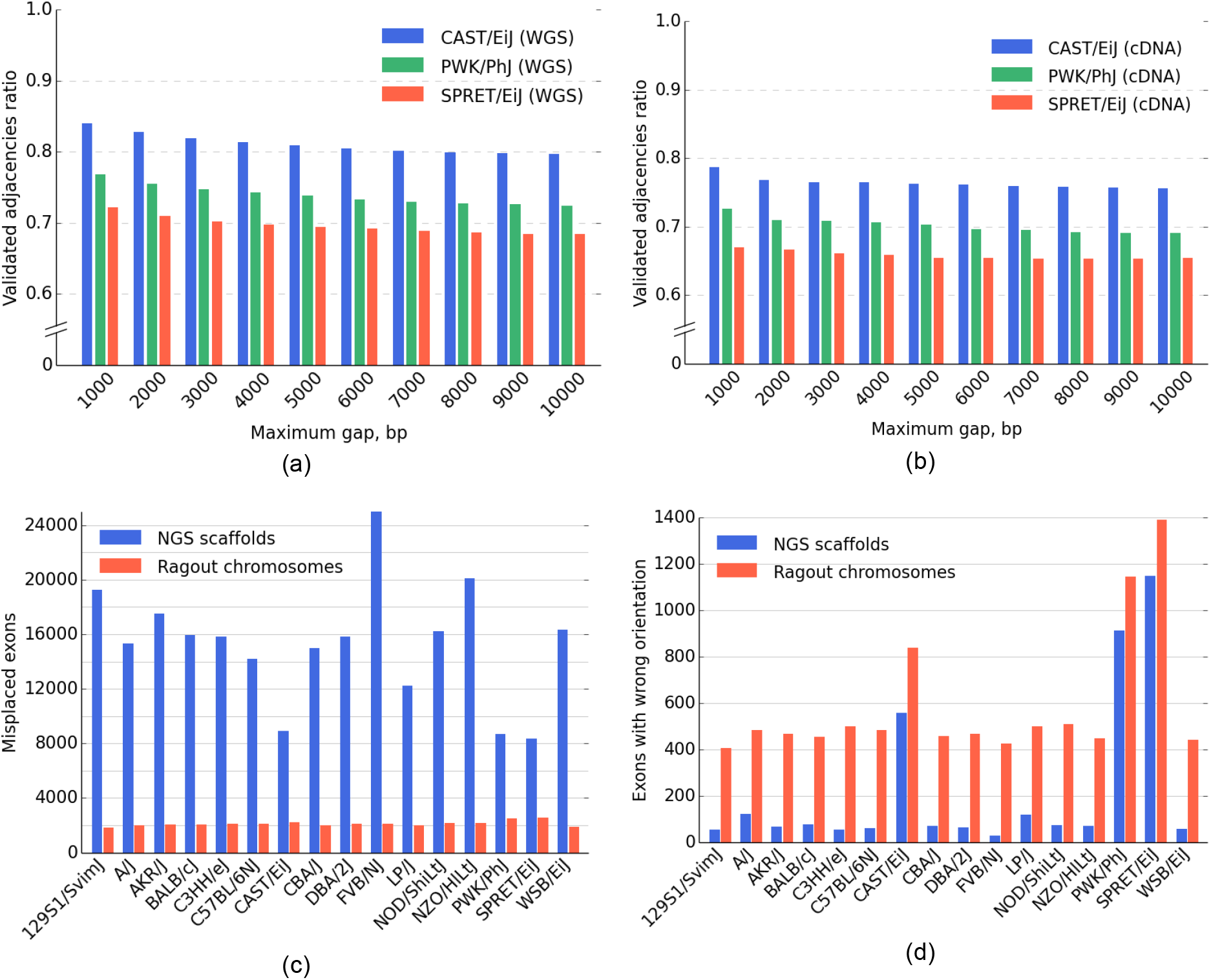
(top) The validated adjacency ratio depending on the maximum gap size of an adjacency for the three most divergent (from *C57BL/6J* reference) *M. musculus* strains. (a) Whole genome sequence data with approximately 0.5x - 1x genome coverage. (b) Whole exome sequencing with approximately 0.3x genome coverage. The probability of a correct and covered adjacency without a gap not being validated by the reads of length 3,000 bp at 1x coverage could be estimated as 15%. (bottom) GENCODE transcript consistency analysis. (c) Number of exons found on non-primary scaffolds/chromosomes. (d) The number of exons on the primary chromosome in a wrong orientation. The total number of transcripts in the database was 78,653. The control analysis of *C57BL/6J* reference genome yielded 1638 misplaced exons and 517 exons in the wrong orientation (due to ambiguous alignments of short exons).

### Mus caroli and Mus pahari assembly

In order to see how our method performs in assembling more distant genomes from the references, we applied Ragout to the *Mus caroli* and *Mus pahari* genomes. These genomes exhibit 4% and 8% sequence divergence from *M. musculus, which is equivalent*, respectively to the human-oragutant and human-marmoset divergence [Thybert et al. 2017]. Since the targets were evolutionarily intermediate between these *M. musculus and Rattus norvegicus*, we used both of these references for the assembly. Additionally, for both *M. pahari* and *M. caroli* assemblies, we used optical maps from the other genome as a third “extra reference” for each genome. The phylogenetic tree of the described four genomes was reconstructed by Ragout.

The *M. caroli* genome was assembled using ALLPATHS-LG [Butler et al., 2008] from NGS short-range libraries, multiple mate-pair libraries with up to 3 kbp insertion length and optical maps. Ragout reconstructed 26 scaffolds (consisting of 28,486 NGS fragments), which correspond to 19 autosomes, chromosome X and 6 small, unlocalized fragments [Thybert et al., 2017]. Detailed assembly statistics are given in Table 2. The assembled *M. caroli* chromosomes do not exhibit any large inter-chromosomal rearrangements with respect to the *M. musculus* genome, which was expected from physical maps as well as chromosome paintings [Thybert et al., 2017]. However, we detected a large cluster of inversions in chromosome 17 of approximately 5 Mbp. In addition to this karyotype-scale variation, we detected 12 synteny block adjacencies that do not appear in any of the three reference genomes, suggesting target-specific rearrangements larger than 10 kbp. In the assembled chromosomes 28 connections between contigs longer than 10 kbp were supported by only the *M. pahari* and/or *R. norvegicus* genomes.

**Table 2.** Statistics for the *M. caroli* and *M. pahari* assemblies using Ragout and RACA. Both tools are able to split input sequences into multiple fragments, thus the sum of the number of used input fragments and unplaced fragments is not necessary equal to the total number of input sequences. Unplaced fragments are defined as the input sequences (or their parts) longer than 1 kb that were not represented in the resulting scaffolds.

| Statistic | *M. caroli* |  | *M. pahari* |  |
| --- | --- | --- | --- | --- |
|  | Ragout | RACA | Ragout | RACA |
| Scaffolds | 26 | 63 | 23 | 56 |
| N50 (kb) | 138,363 | 86,701 | 145,528 | 95,590 |
| Assembly length (mb) | 2,556 | 2,476 | 2,477 | 2,454 |
| Used NGS fragments | 27,642 | 9,679 | 15,349 | 6,748 |
| Unplaced NGS fragments | 5,792 | 20,291 | 4,159 | 10,726 |
| Unplaced length (kb) | 55,865 | 344,887 | 49,747 | 181,150 |

We also applied RACA to assemble *M. caroli* using the same set of references and all available sequencing libraries (see Table 2). Synteny block of size was set to 50 kb as it previously produced the optimal results on *M. musculus* genomes in terms of assembly contiguity and coverage. Some of the assembled chromosomes were left fragmented (63 scaffolds, N50 = 86 Mbp) and a significant portion of sequence (344 Mbp) was left unplaced by RACA, which could be explained by the fact that only a single fixed synteny block size is used. In contrast, an assembly with synteny block scale of 10 kb left 63 Mb of sequence unplaced, but N50 was two times lower (40 Mbp).

The same protocol was used to assemble the *M. pahari* genome (see the results in Table 2). Ragout reconstructed 23 scaffolds from 16,108 assembly fragments. In contrast to *M. caroli*, chromosome painting as well as physical mappings of *M. pahari* suggest extensive inter-chromosomal rearrangements [Thybert et al., 2017]. Ragout detected five chromosome fusions, four of which are consistent with the mappings and one that is supported by the *R. norvegicus* reference and might have been missed from the optical maps due to its relatively small size (about 2 Mbp). The dot-plots of the detected large-scale rearrangements as well as their corresponding chromosome paintings are shown on Figure 4. Four expected chromosome fusions and 13 expected chromosome fissions were not detected by our algorithm. This was mainly caused by the missing signatures of the rearrangements in the input sequences; in particular, it is currently very hard to predict a chromosome fission if it is not supported by any of the references, since target fragments can not provide positive evidence of such an event. Ragout resolves this issue by integrating physical mappings. Importantly, Ragout did not generate any large inter-chromosomal rearrangements which were not expected from physical mappings or references. We also detected 36 adjacencies that do not appear in any of the reference genomes, suggesting target-specific rearrangements of size more than 10 kbp. Twenty one connections between contigs longer than 10 kbp in the final chromosomes were supported by only the *M. caroli* and/or *R. norvegicus* genomes. On the same dataset RACA produced an assembly with a higher number of scaffolds and lower N50 (see Table 2). RACA found four of the five chromosome fusions detected by Ragout.

We also detected a cluster of inversions with the same structure as in the *M. caroli* genome in a chromosome, homologous to a *M. musculus* chromosome 17. The breakpoints of the detected inversions in both genomes were supported by the corresponding optical maps. Interestingly, both the *M. musculus* and *R. norvegicus* references contain different structural variations within this region and share one inversion breakpoint (see Fig. 5). This might be a signature of a rearrangement hotspot [Pevzner et al., 2003].

## Discussion

Despite recent advances in sequencing technologies and bioinformatics algorithms, *de novo* assembly of a mammalian scale genome into a complete set of chromosomes remains a challenge. Many genome sequencing projects [Wang et al., 2014, Dobrynin et al., 2015, Vij et al., 2016] have used reference-guided assembly as a step in genome finishing, often followed by manual sequence curation. While multiple tools for reference-assisted assembly exist, their performance has proved limited when the reference genomes exhibit a significant number of structural variations relative to the target genome being assembled. In this manuscript we have presented Ragout - an algorithm for chromosome assembly of large and complex genomes using multiple references. Ragout joins input NGS contigs or scaffolds into larger sequences by analyzing genome rearrangements between multiple references and the target genome. In difference to previous approaches, Ragout utilizes hierarchical synteny information, which helps to reduce gaps in the resulting chromosomes. We used simulations to show that Ragout makes few errors, even in the presence of complicated rearrangements, and outperforms previous approaches in both accuracy and assembly completeness.

Using the existing *Mus musculus* reference, we applied Ragout to assemble 15 *Mus musculus* and one *Mus spretus* genomes (including 12 laboratory strains and four wild-derived strains). Through the benchmarks, which included validations with long PacBio reads, transcriptome analysis and PCR testing, we show that Ragout produced highly accurate chromosome assemblies with less than 5% of sequence unplaced (2% of coding sequence). An analysis of transcript data showed a substantial improvement of the resulting assemblies from a gene structure perspective. Importantly, our algorithm is capable of preserving target-specific rearrangements, which are observed in the NGS assembly but not in the reference genomes, thus having less reference bias than simpler reference-guided approaches. Using both the reference *M. musculus* and *R. norvegicus* reference genomes, we used Ragout to assemble two more challenging genomes: *M. pahari* and *M. caroli*, which exhibit major karyotype-scale differences compared to these references. *M. caroli* was assembled into a complete set of chromosomes with the expected karyotype structure. *M. pahari* chromosomes assembled by Ragout contained five large intra-chromosomal rearrangements, four of which were confirmed by chromosome painting techniques. The fifth rearrangement was supported by the *R. norvegicus* reference and might have been missed from chromosome maps due to its relatively small size. While Ragout did not generate any unexpected rearrangements (not supported by chromosome maps or references), some expected rearrangements remained undiscovered. Most of these rearrangements were chromosome fissions, which are difficult to predict using only NGS sequencing data. Ragout has a module to incorporate chromosome maps to guide the final assembly and successfully incorporated all expected rearrangements into the final chromosomes.

One limitation of Ragout is that the algorithm currently does not support diploid genomes, thus only a single copy of each chromosome is reconstructed (possibly, as a mixture of different alleles). Complete *de novo* diploid genome assembly remains a challenging task, even with recent improvements in sequencing technologies, which include long PacBio reads or long Hi-C interactions that can link heterozygous variations. Being mostly orthogonal problems, the information from reference genomes could be further coupled with the direct sequencing approaches to improve *de novo* diploid genome assembly and phasing.

While Ragout uses 2-break rearrangement model to distinguish chimeric adjacencies from real rearrangements, the RACA algorithm implements a different approach, in which the information from the paired-end sequencing libraries is used detect unreliable scaffold connections. The two approaches are orthogonal and ideally should be combined together in the future to get the optimal assembly results.

We have shown that Ragout produces accurate and complete chromosome assemblies of mammalian-scale genome even in the presence of extensive structural rearrangement. The assembled chromosomes were used as starting points for manual sequence curation in the mouse strains assembly project [Keane et al., in preparation, Thybert et al., 2017].

### Availability

The complete Ragout package is freely available at: http://fenderglass.github.io/Ragout/. The genomes of 15 *Mus musculus* and *Mus spretus, which* have been scaffolded using Ragout with additional curations, are available at NCBI database under *PRJNA310854* bio project id [Keane et al., in preparation]. The genome assemblies of *M. caroli* and *M. pahari* were submitted to the European Nucleotide Archive (www.ebi.ac.uk/ena) and are available with accession numbers GCA_900094665 (*M. caroli*) and GCA_900095145 (*M. pahari*).

**Figure 4.**
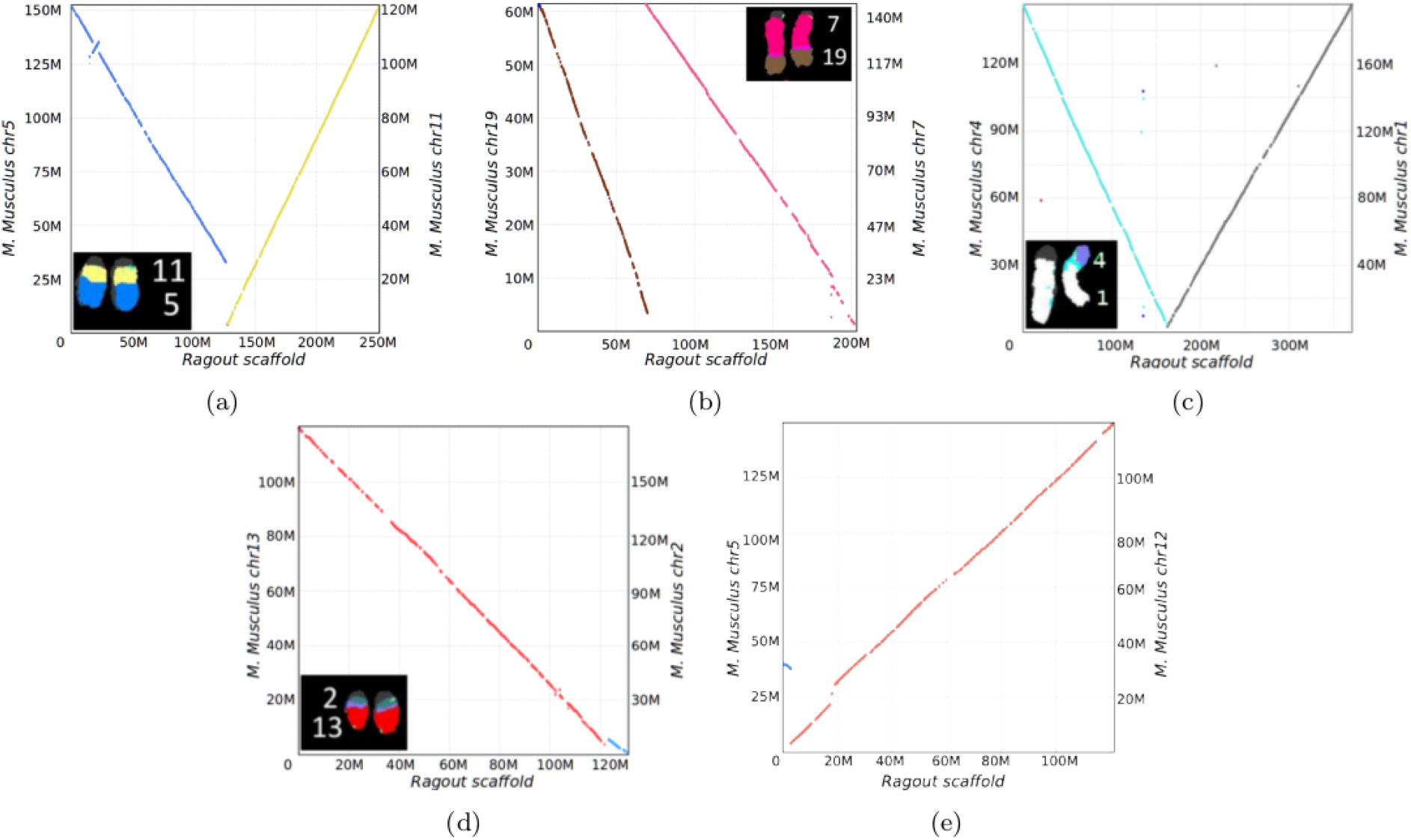
Dot-plots and corresponding chromosome paintings showing inter-chromosomal rearrangements in *M. pahari* assembly. The rearrangement in (e) is supported by the *R. norvegicus* reference and might be missed from chromosome coloring due to its small size (about 2 Mbp).

**Figure 5.**
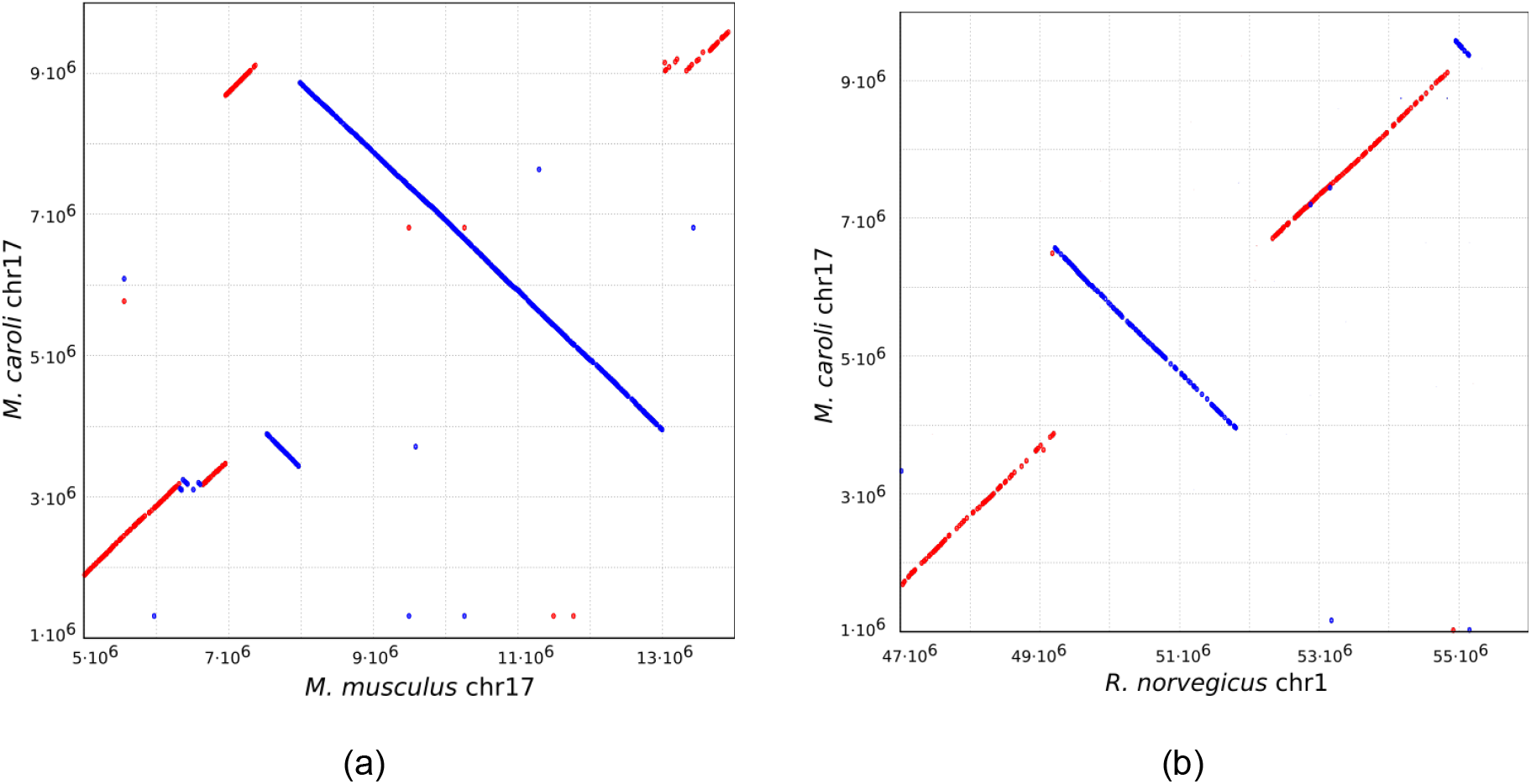
Dot-plots of a chromosome 17 region of *M. caroli* against (a) *M. musculus* and (b) *R. norvegicus* showing a cluster of structural variations of size approximately 5 Mbp. *M. pahari* chromosomes shows the same genomic structure. The inversion breakpoints are supported by the optical maps in both *M. caroli* and *M. pahari* genomes. *M. musculus* and *R. norvegicus* references are sharing one breakpoint of two different inversions.

## Methods

### Construction of synteny blocks

Ragout constructs synteny blocks at multiple scales and provides a hierarchical relationship among these different scales of synteny blocks (for a complete discussion of multi-scale synteny blocks, see [Minkin et al., 2013, Ghiurcuta and Moret, 2014, Kolmogorov et al., 2014]). At the highest level of resolution, the alignment of multiple genomes is represented as a set of alignment blocks. Each alignment block is a set of oriented, non-overlapping, homologous sub-intervals of the input genomes. To derive a set of alignment blocks we use Progressive Cactus. Each genomic sequence *s_t_* can be represented in the alphabet of alignment blocks *s_t_ = b_1_, …, b_n_*, where *b_i_* is an alignment block of length *|b_i_|*. Given a set of *k* sequences *S* = {*s_1_, …, s_k_*} and a parameter *minBlock*, we construct an A-Bruijn graph *G(S, minBlock)* as follows: For each alignment block b_i_ such that |b_i_| ≥ minBlock we create two nodes *b^h^_i_* and *b^t^_i_* (representing the head and tail of each block) and connect them by a black edge. We connect heads and tails of adjacent alignment blocks in each genome s_t_ with an adjacency edge with color *C_t_*. The length of an adjacency edge is defined as the distance between the two alignment blocks in the corresponding genome. We also include a special infinity node [Alekseyev and Pevzner, 2009] representing the ends of sequences (telomeres in complete genomes or fragment ends in draft genomes). A node is called a bifurcation if it is connected with more than one node (by colored edges) or is connected with an infinity node. A path without any bifurcation node except for its start and end is called non-branching. Figure 6b shows an example of an A-Bruijn graph constructed from alignment blocks present in Figure 6a. Intuitively, this construction is equivalent to: (1) representing each sequence in the alphabet of alignment blocks and (2) gluing together two alignment blocks if they are homologous and have size larger than *minBlock* (See [Medvedev et al., 2011] for a formal definition of gluing). Constructing this A-Bruijn graph is also equivalent to constructing the breakpoint graph from multiple genomes (See [Lin et al., 2014] for a proof of equivalence).

Small polymorphisms and micro-rearrangements correspond to certain cycle types or local structures in the A-Bruijn graph and disrupt large homologous regions into many shorter ones (see [Pham and Pevzner, 2010] for a thorough cycle type classification of an A-Bruijn graph). We use a progressive simplification algorithm to simplify bulges and parallel paths in the A-Bruijn graph. The algorithm consists of two sub-procedures: *CompressPaths* and *CollapseBulges*, and is parameterized with a value *maxGap*, which is the maximum length of the cycle/path to be simplified. *CompressPaths* is used to merge collinear alignment blocks into a larger synteny block. It starts from a random bifurcation node and traverses the graph in an arbitrary direction until it reaches another bifurcation node or an adjacency edge that is longer than *maxGap*. Then, it exchanges the traversed path with two nodes connected by a black edge (which corresponds to a new synteny block, see Figure 6e for an example). A complementary procedure – *Collapse-Bulges* – finds all simple bulges having branch length shorter than *maxGap* in a similar manner and exchanges the bulge with a single synteny block as described previously (see Figure 6b). These two procedures are applied one after another multiple times, until no more simplification can be done. It is easily verified that the result is invariant to the order of such operations. After the simplification step, the sequences of synteny blocks are recovered by “threading” the genomes through the graph. If the blocks require additional simplification, a larger value of *minBlock* is used to construct the new A-Bruijn graph (see Figure 6d). See *Supplementary Materials S3* for an extra discussion on the choice of *maxGap* and *min-Block*.

**Figure 6.**
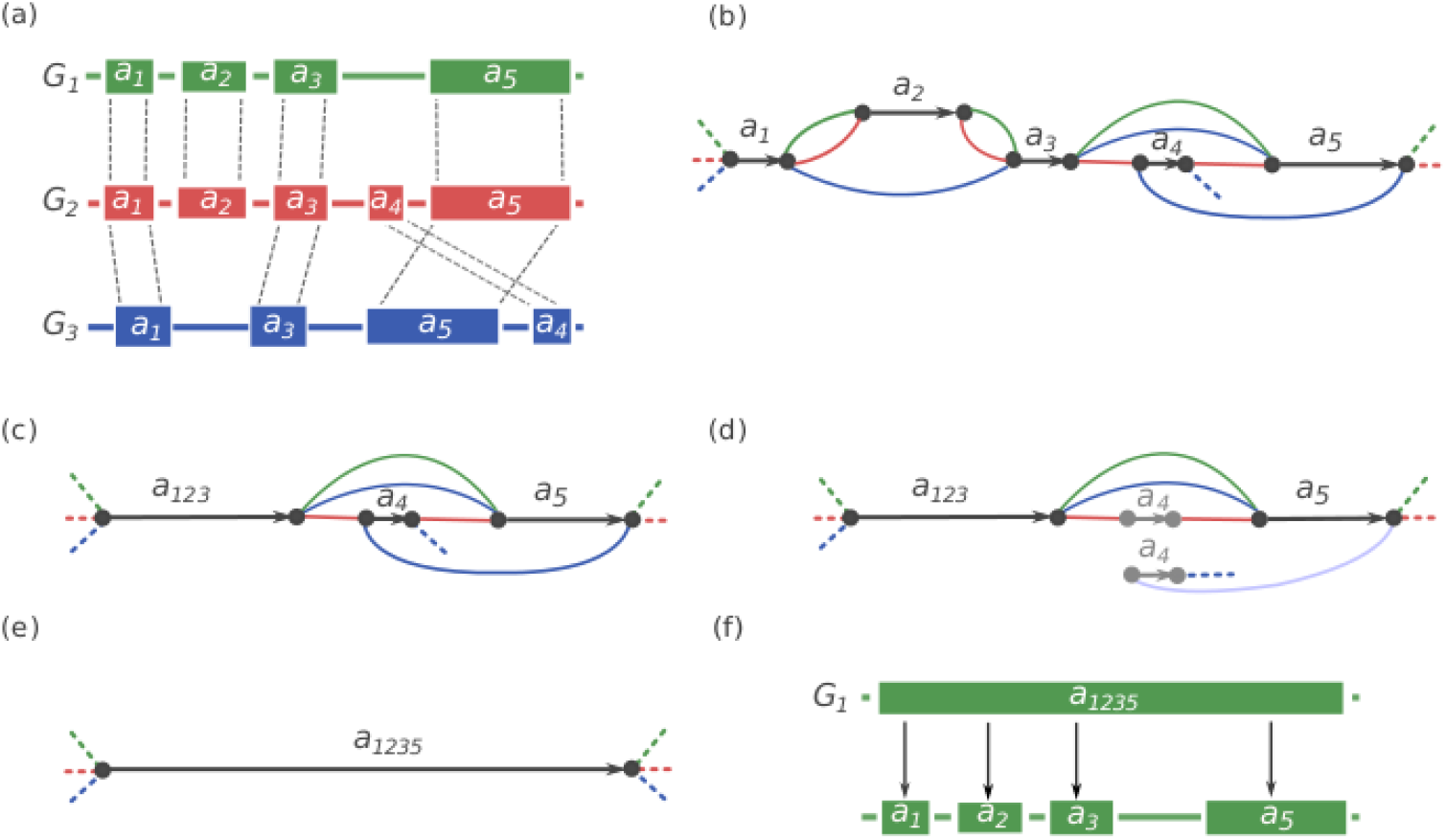
Synteny blocks reconstruction. (a) Three genomes *G_1_, G_2_, G_3_* are encoded in the alphabet of alignment blocks. *a_1_, a_2_, a_3_, a_4_, a_5_. |a_4_| < |a_1_| < |a_3_| < |a_2_| < |a_5_|*. (b) The A-Bruijn graph constructed with *minBlock = |a_4_|*. (c) The A-Bruijn graph after bubble simplification. *a_123_* represents the new merged block from *a_1_, a_2_, a_3_*. (d) A next iteration of the A-Bruijn graph constructed with *|a_4_| < minBlock* ≤ |a*_1_|*, which eliminates block *a_4_*. (e) *a_123_*, and *a_5_* are merged into a larger block *a_1235_*. (f) The hierarchical representation of synteny blocks: the larger block *a_1235_* can be decomposed into smaller blocks *a_1_, a_2_, a_3_* and *a_5_*.

### Incomplete breakpoint graphs and missing adjacency inference

After running the synteny procedure, the input genome sequences can be represented in the alphabet of synteny blocks. For the sake of simplicity, let us assume that every synteny block is represented exactly once in each genome and all reference genomes are complete (the issue of repetitive synteny blocks will be addressed further in the manuscript). Given an assembly *A* and *k* reference sequences *P_1_*, …, *P_k_* in the alphabet of synteny blocks *B*, we construct the incomplete multi-color breakpoint graph *BG*(*A*, *P_1_*, …, *P_k_*) = (*V*, *E*), where *V = {b^h^i, b^t^i |b_i_* ∈ B} [Kolmogorov et al., 2014]. For each synteny block, there are two vertices in the graph which correspond to the tail and head of the block. Edges are undirected and colored by *k + 1* colors. An edge connects vertices that correspond to heads/tails of adjacent synteny blocks and is colored by the corresponding color of the genome/assembly (Figure 1). We use red, *P_1_*, …, *P_k_* to refer to the colors of edges, where red edges represent the adjacencies of synteny blocks in the target assembly *A*, and *P_i_* represents the adjacencies of synteny blocks in genome *P_i_*. If the target genome were available, the set of all red edges would define a perfect matching in the graph. However, since the genome is fragmented into contigs, the adjacency information of the target genome at the vertices that correspond to the end of contigs is missing. Ragout infers these missing red edges in the graph by solving the half-breakpoint parsimony problem using other adjacencies from the reference genomes as well as using their evolutionary relationship in the form of a phylogenetic tree (See [Kolmogorov et al., 2014] for a complete description of the half-breakpoint parsimony problem). In contrast to bacterial assemblies, mammalian assemblies usually contain a higher fraction of chimeric contigs/scaffolds due to increased size and complexity. These problematic contigs generate many erroneous red connections in the breakpoint graph and need to be removed before inferring the missing adjacencies. Below, we describe a new algorithm that uses rearrangement signatures in the breakpoint graph to distinguish rearrangement red edges from erroneous red edges.

### Detection of chimeric sequences

Mammalian assemblies usually contain high numbers of misassembled contigs. A large portion of misassembled contigs are chimerics, which are when an NGS assembler artificially joins regions in the assembled genome that are not truly adjacent. These false adjacencies correspond to erroneous red edges in the breakpoint graph. Erroneous red edges should be identified and removed before the missing adjacency inference step, since misassemblies in different contigs will join together and cause large structural errors. We denote a red edge in the breakpoint graph as *supported*, if there is at least one parallel reference edge, otherwise the red edge is denoted *unsupported*. We call an adjacency *genomic* if it is a true adjacency present in the target genome and *artificial* if it is a result of a chimeric contig. Supported red edges are unlikely to be artificial; however, an unsupported red edge could be either genomic (coming from a rearrangement specific to the target genome) or artificial (in case of misassemblies). To classify each red edge as either genomic or artificial, we need to analyze the rearrangement structure between the target genome and references. Since any rearrangement operation can be modeled as a k-break operation [Alekseyev and Pevzner, 2009], rearrangement will generate alternating cycles in the breakpoint graph (see Figure 8 for an example). For each unsupported edge, if it does not belong to an alternating cycle, we classify it as artificial and remove it from the breakpoint graph.

**Figure 7.**
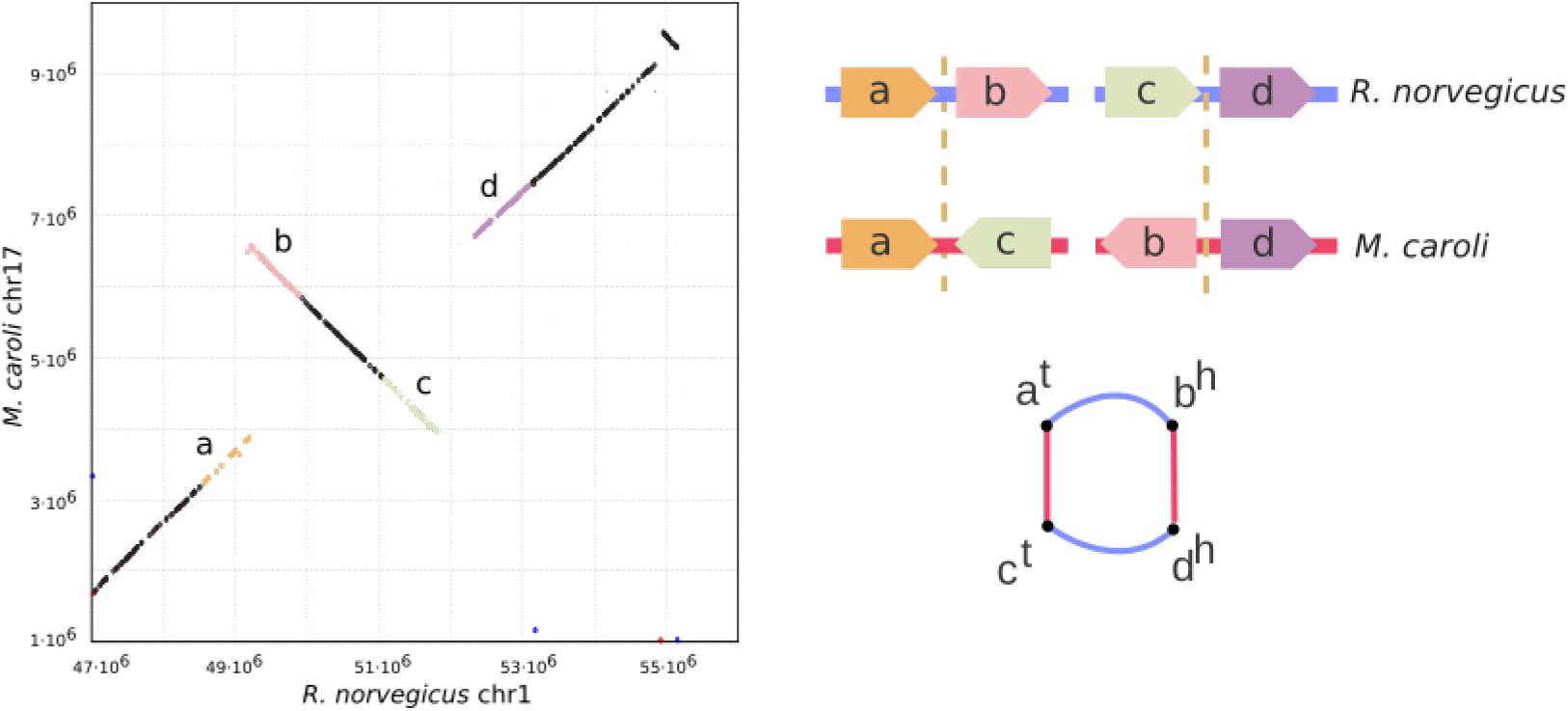
An inversion recovered by Ragout forms an alternating cycle of length four in the breakpoint graph. Both red edges are classified as supported.

### Iterative assembly

We first perform multiple rounds of scaffold assembly with different synteny block resolution. The scaffolds from the largest scale represent a “skeleton” for the final assembly. We then iteratively merge the existing skeleton with the set of scaffolds constructed from finer-scale synteny blocks, so as the the new assembly is consistent with the skeleton structure (see Kolmogorov et al., [2014] for the details). By default, Ragout is run in three iterations with synteny blocks sizes equal to (10000, 500, 100), which are similar to the default synteny scales in Sibelia, but with an increased block size in the first iteration to account for longer and more complex repeats in mammalian genomes. See additional details of the merging algorithm in *Supplementary Materials S4*.

### Repeat resolution

The breakpoint graph analysis requires all synteny blocks to appear exactly once in each genome. A common approach is to filter out all repetitive blocks before analyzing the breakpoint graph. While this approach works for bacterial genomes, it is not applicable for mammalian assemblies, since the number of unresolved repeats in mammalian assemblies is much larger. Filtering out all such sequences would lead to a large number of gaps in the final assembly. Here we present an algorithm that addresses this issue by resolving repeat sequences based on the information from the references. Intuitively, for each unresolved repeat, we want to create *k* different instances of that repeat, where *k* is the copy number of the repeat in the target genome. As some repeats are already resolved by the NGS scaffolder, the corresponding synteny blocks in contigs will be surrounded by other unique synteny blocks. We use this “context” information to map repeat instances in contigs to the corresponding repeats in reference genomes. See additional details about repeat resolution algorithm in *Supplementary materials S5*.

### Phylogenetic tree reconstruction

When the phylogenetic tree of the input sequence is not available, it can be inferred from the adjacency information [Lin et al., 2011]. Given a set of genomes represented as permutations of synteny blocks, for each pair of genomes we first build a breakpoint graph. Let *b_1_* be a set of all breakpoints (graph edges) from the first genome and *b_2_* from the second genome. We denote *b_1_ & b_2_* as a set of breakpoints that are shared in both genomes. The distance between two genomes is defined as *min{size(b_1_), size(b_2_*)} – size(b*_1_ & b_2_)*. We then build a distance matrix and run standard Neighbor-Joining algorithm [Saitou and Nei, 1987], which provides a good approximation of a real phylogenetic tree [Moret et al., 2001].

## Supplementary Materials

### S1 Benchmarking RACA on *Mus Musculus* and *Mus Spretus* genomes

We compared Ragout performance against RACA using three *Mus* genomes for which long PacBio reads were available (*PWK/PhJ, SPRET/EiJ and CAST/EiJ)*. Similarly to Ragout, RACA runs were performed using one reference (*C57BL/6J*) as well as all available paired-end and mate-pair libraries. To select an optimal synteny block resolution, each genome was assembled multiple times with different minimum block size threshold: 10 kb, 50 kb and 150 kb (as in the original manuscript [Kim et al, 2013]). The detailed statistics are given in Supplementary Table 1. The assemblies produced by RACA had the expected chromosome structure for the larger block sizes (50kb and 150kb), but the assemblies with the block size of 10kb were more fragmented. As expected, the total length of unplaced sequence was also positively correlated with increase of the synteny blocks scale. Interestingly, all RACA assemblies had more unplaced sequence and lost exons than the corresponding Ragout assemblies, which highlights the benefits of iteraretive assembly with multiple synteny block scales. We then used the GENCODE gencode transcript set to compare the structural accuracy of Ragout and RACA chromosomes (see Supplementary Table 1). Ragout chromosomes consistently showed less split transcripts and transcripts with wrong orientation, while the number PacBio reads with correct alignment orientations was higher.

**Supplementary Table 1.** Statistics for RACA assemblies of *PWK/PhJ, SPRET/EiJ and CAST/EiJ* with different synteny block scale.

| Dataset | Scaffolds | Scaffolds length (mb) | N50 (mb) | Unplaced (mb) | Lost exons |
| --- | --- | --- | --- | --- | --- |
| *CAST/EiJ*, 10 kb | 37 | 2,552 | 115 | 141 | 8,650 |
| *CAST/EiJ*, 50kb | 20 | 2,544 | 131 | 150 | 9,707 |
| *CAST/EiJ*, 150kb | 21 | 2,536 | 128 | 158 | 10,383 |
| *PWK/PhJ*, 10kb | 48 | 2,454 | 95 | 129 | 9,742 |
| *PWK/PhJ*, 50kb | 22 | 2,360 | 124 | 223 | 21,143 |
| *PWK/PhJ*, 150kb | 19 | 2,356 | 128 | 227 | 21,402 |
| *SPRET/EiJ*, 10kb | 44 | 2,536 | 97 | 132 | 8,080 |
| *SPRET/EiJ*, 50kb | 23 | 2,536 | 128 | 132 | 7,968 |
| *SPRET/EiJ*, 150kb | 20 | 2,536 | 130 | 132 | 8,538 |

**Supplementary Table 2.** Structural accuracy comparison of RACA and Ragout chromosomes. All statistics are given as differences between the corresponding values calculated for RACA and Ragout results. Consistent PacBio reads were defined as reads that have all local alignments with the same orientation.

| Dataset | Missplaced exons,<br>#RACA - #Ragout | Exons with wrong orientation,<br>#RACA - #Ragout | Consistent PacBio reads,<br>#Ragout - #RACA |
| --- | --- | --- | --- |
| *CAST/EiJ*, 10 kb | 1,457 | 484 | 3,386 |
| *CAST/EiJ*, 50kb | 1,211 | 470 | 4,225 |
| *CAST/EiJ*, 150kb | 1,460 | 529 | 2,196 |
| *PWK/PhJ*, 10kb | 1,762 | 494 | 1,494 |
| *PWK/PhJ*, 50kb | 1,441 | 627 | 6,127 |
| *PWK/PhJ*, 150kb | 1,359 | 625 | 6,145 |
| *SPRET/EiJ*, 10kb | 1,559 | 423 | 1,809 |
| *SPRET/EiJ*, 50kb | 1,553 | 609 | 1,444 |
| *SPRET/EiJ*, 150kb | 1,489 | 568 | 934 |

### S2 Benchmarking Ragout and RACA on multiple human genome assemblies

We benchmarked the performance of the Ragout and RACA algorithms using the multiple human chromosome 14 assemblies that were previously generated by different NGS assemblers as a part of the GAGE assembly evaluation study [Salzberg et al., 2012]. The scaffolding results for RACA were taken from the original manuscript [Kim et al, 2013]. Similarly to the previous benchmarks, we selected 50 kb synteny block resolution as the most optimal for all assemblies except SGA, for which 10 kb was chosen. Similarly, we used *M. musculus* and *P. abelii* references to assemble the NGS contigs into chromosomes using Ragout. We evaluated accuracy of the assemblies by aligning them on the reference chromosome using LASTZ [Harris, 2007] and calculating the number of breakpoints between alignments with inconsistent order / orientation. The misassembly rates were computed on two scales: 5 kb and 50 kb. The results are ginven in the Supplementary Table 3. The error rates for both tools were highly correlated accross the datasets, confirming that the choice of initial contigs and a reference genome is critical for the reference chromosome assembly. For most of the datasets Ragout performed equally or better than RACA in terms of N50 and structurall accuracy.

**Supplementary Table 3.**
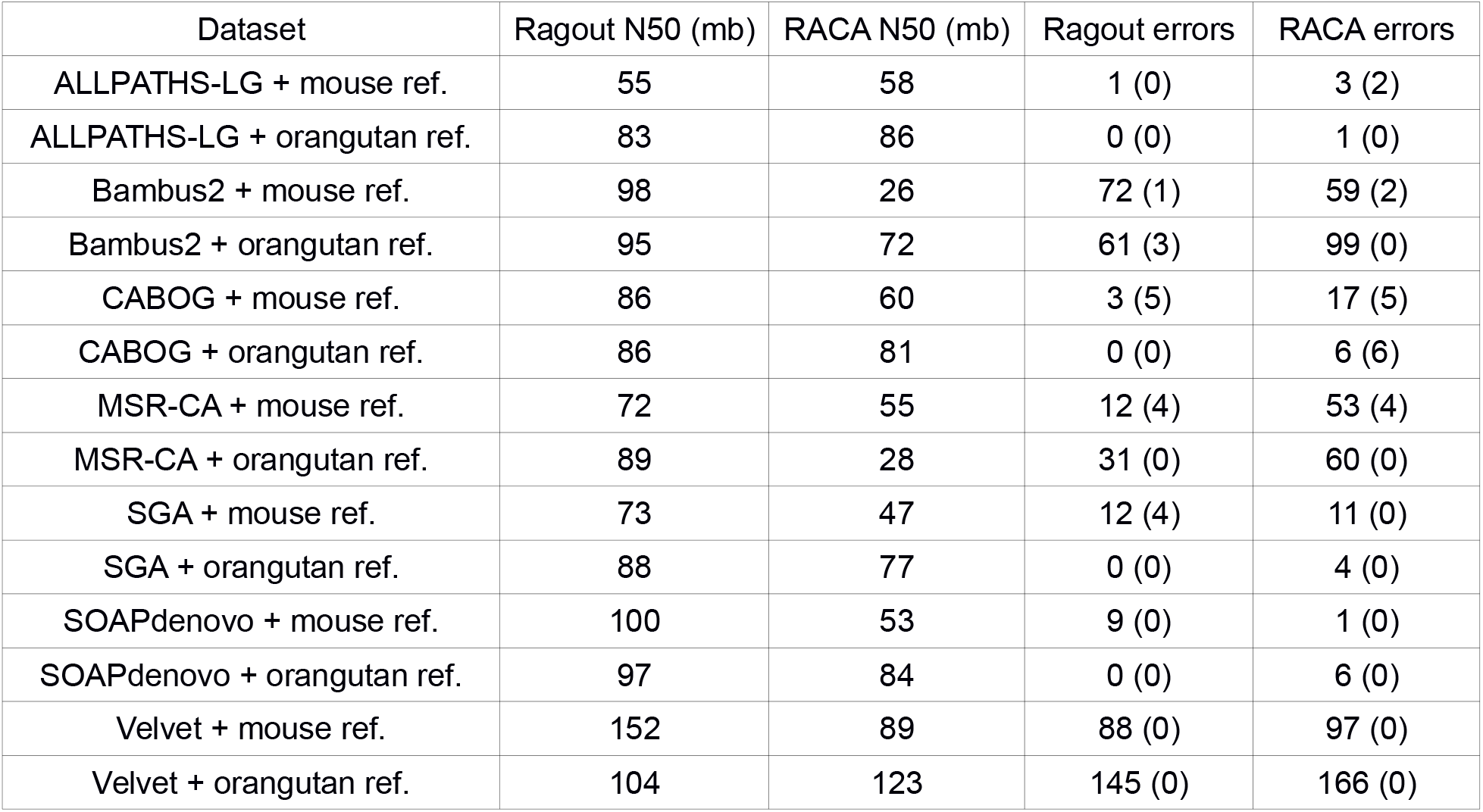
Comparison of Ragout and RACA using the multiple human chromosome 14 assemblies from the GAGE dataset. The misassembly errors were computed for alignments of size more than 5 kb (50 kb).

| Dataset | Ragout N50 (mb) | RACA N50 (mb) | Ragout errors | RACA errors |
| --- | --- | --- | --- | --- |
| ALLPATHS-LG + mouse ref. | 55 | 58 | 1 (0) | 3 (2) |
| ALLPATHS-LG + orangutan ref. | 83 | 86 | 0 (0) | 1 (0) |
| Bambus2 + mouse ref. | 98 | 26 | 72 (1) | 59 (2) |
| Bambus2 + orangutan ref. | 95 | 72 | 61 (3) | 99 (0) |
| CABOG + mouse ref. | 86 | 60 | 3 (5) | 17 (5) |
| CABOG + orangutan ref. | 86 | 81 | 0 (0) | 6 (6) |
| MSR-CA + mouse ref. | 72 | 55 | 12 (4) | 53 (4) |
| MSR-CA + orangutan ref. | 89 | 28 | 31 (0) | 60 (0) |
| SGA + mouse ref. | 73 | 47 | 12 (4) | 11 (0) |
| SGA + orangutan ref. | 88 | 77 | 0 (0) | 4 (0) |
| SOAPdenovo + mouse ref. | 100 | 53 | 9 (0) | 1 (0) |
| SOAPdenovo + orangutan ref. | 97 | 84 | 0 (0) | 6 (0) |
| Velvet + mouse ref. | 152 | 89 | 88 (0) | 97 (0) |
| Velvet + orangutan ref. | 104 | 123 | 145 (0) | 166 (0) |

### S3. The choice of parameters for A-Bruijn graph simplification

The choices of *minBlock* and *maxGap* are not independent – their scale should be in agreement with each other. Using a small value of minBlock with a big value of *maxGap* will lead to longer blocks with low similarity (false blocks). On the other hand, if *maxGap* is much smaller than *minBlock*, the effect of simplification will be minor. Moreover, we cannot start from big *minBlock* as the initial alignment might be highly fragmented into many small alignment blocks. As a solution for these issues, we run the simplification algorithm iteratively. Starting from small *minBlock* and *maxGap*, we gradually increase them until we reach the target synteny block scale. The values of *minBlock* and *maxGap* are chosen empirically by comparing the final results with 2-D synteny dot-plot pictures. For a complete discussion about multi-scale synteny blocks, see [Minkin et al., 2013, Ghiurcuta and Moret, 2014, Kolmogorov et al., 2014].

### S4. Additional details on the iterative assembly algorithm

Let *S_skeleton_* be a current skeleton of scaffolds and *S_new_* the scaffold set that is being merged. The merging algorithm consists of two parts: rearrangement projection and gap filling. During the rearrangement projection we first detect rearrangements between *S_new_* and *S_skeleton_* by constructing a 2-color breakpoint graph. Nodes in this graph correspond to the contigs in *S_skeleton_* and *S_new_*. As some target-specific rearrangements were not detected in *S_skeleton_*, but appear in *S_new_*, they will form alternating cycles on the breakpoint graph. We then apply (project) the newly detected rearrangements to the *S_skeleton_*. Not all rearrangements can be safely projected, because some of them might be erroneous (since smaller synteny blocks are less reliable). We call a rearrangement safe, if it (i) involves fewer than *k* breaks, and (ii) the chromosomal similarity before and after applying this rearrangement to *S_skeleton_* is more than *c*. Chromosomal similarity is defined as the percentage of synteny blocks that stay in the same scaffold after applying the rearrangement. This prevents large chromosomal translocations, fusions and fissions from being projected. We found that *k = 4* and *c = 0.9* work well in most cases. This setting also allows all common rearrangement types: inversion, transposition, small chromosomal translocation and gene conversion. After projecting rearrangements, we insert small contigs from *S_new_* to *S_skeleton_*, such as the resulting contig order is consistent with the order in S_skeleton_ as described in [Kolmogorov et al., 2014].

### S5. Additional details for the repeat resolution algorithm

Repetitive synteny blocks are defined as blocks with at least two copies in a genome. We denote a set of all instances of a single repetitive synteny block inside a genome as a repeat family. Given a repeat family *RF*, for each instance in *RF* we define context as an ordered set of at most *2b* closest synteny blocks (*b* from the left and *b* from the right, any or all of which may be repetitive as well). We call a repetitive block resolved, if there is at least one unique synteny block among these *2b* blocks. If all blocks in the context are repetitive or the context is empty, the repetitive block is called unresolved. It is expected that orthologous copies of a repeat share a similar context during genome evolution. Given two contexts *c_1_* and *c_2_*, we define *score(c_1_, c_2_)* as the alignment score between the sequences of blocks from *c_1_* and *c_2_* (a match between unique/repetitive blocks adds +2/+1 to the score, respectively; mismatches and gaps are penalized by -1). Let *G_r_* and *G_t_* be the reference and the target genomes, respectively. First, we find a mapping between *R_r_* and *R_t_*, which are resolved blocks from G_r_ and G_t_, respectively. We construct a full bipartite graph, where nodes from each set correspond to repetitive blocks from *R_r_* and *R_t_*, respectively. Then for each pair of nodes *r_i_*, *r_j_* from two sets, we put an edge with weight equal to *score(c_i_, c_j_)*, where *c_i_* and *c_j_* are contexts of *r_i_*, *r_j_*. We then find a maximum weight matching, which corresponds to an optimal mapping between *R_r_* and *R_t_*.

After finding the matching for resolved blocks, we apply a similar approach for unresolved ones, with an exception that one repeat block from the target assembly may match against multiple repeats from the reference. The above algorithm can be extended in the case of multiple references by using a strategy similar to the progressive method for multiple sequence alignment.

### S6. Additional information on simulated dataset

**Supplementary Table 4.**
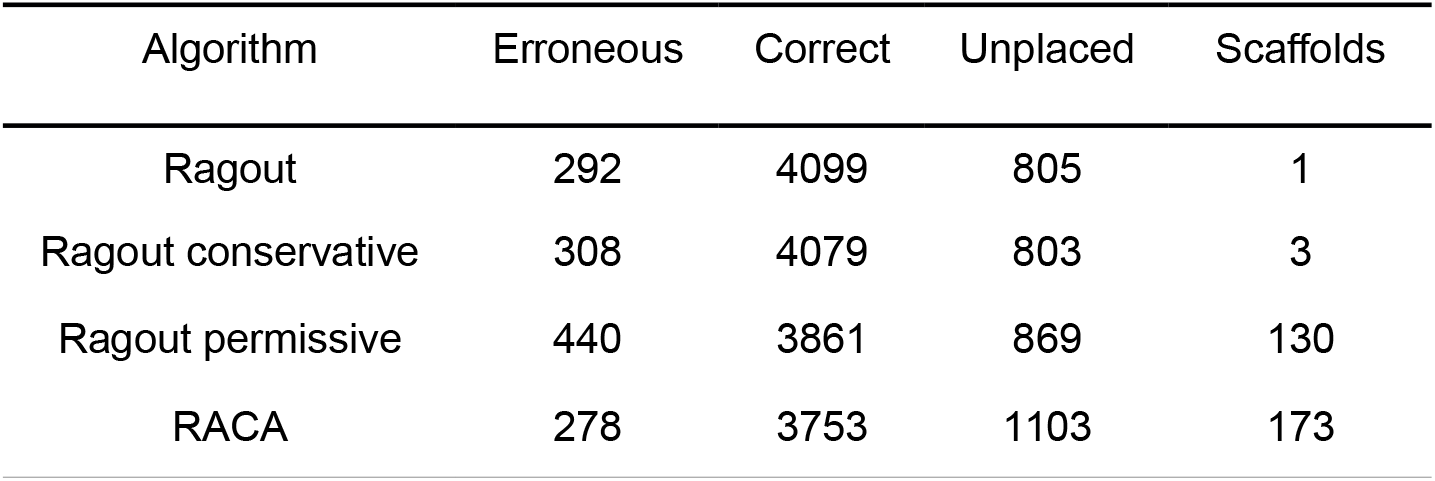
Detailed assembly statistics for a simulated dataset with 50 rearrangements. Assembly consists from 4,926 fragments with a mean length 18,000 bp.

| Algorithm | Erroneous | Correct | Unplaced | Scaffolds |
| --- | --- | --- | --- | --- |
| Ragout | 292 | 4099 | 805 | 1 |
| Ragout conservative | 308 | 4079 | 803 | 3 |
| Ragout permissive | 440 | 3861 | 869 | 130 |
| RACA | 278 | 3753 | 1103 | 173 |

**Supplementary Figure 1.**
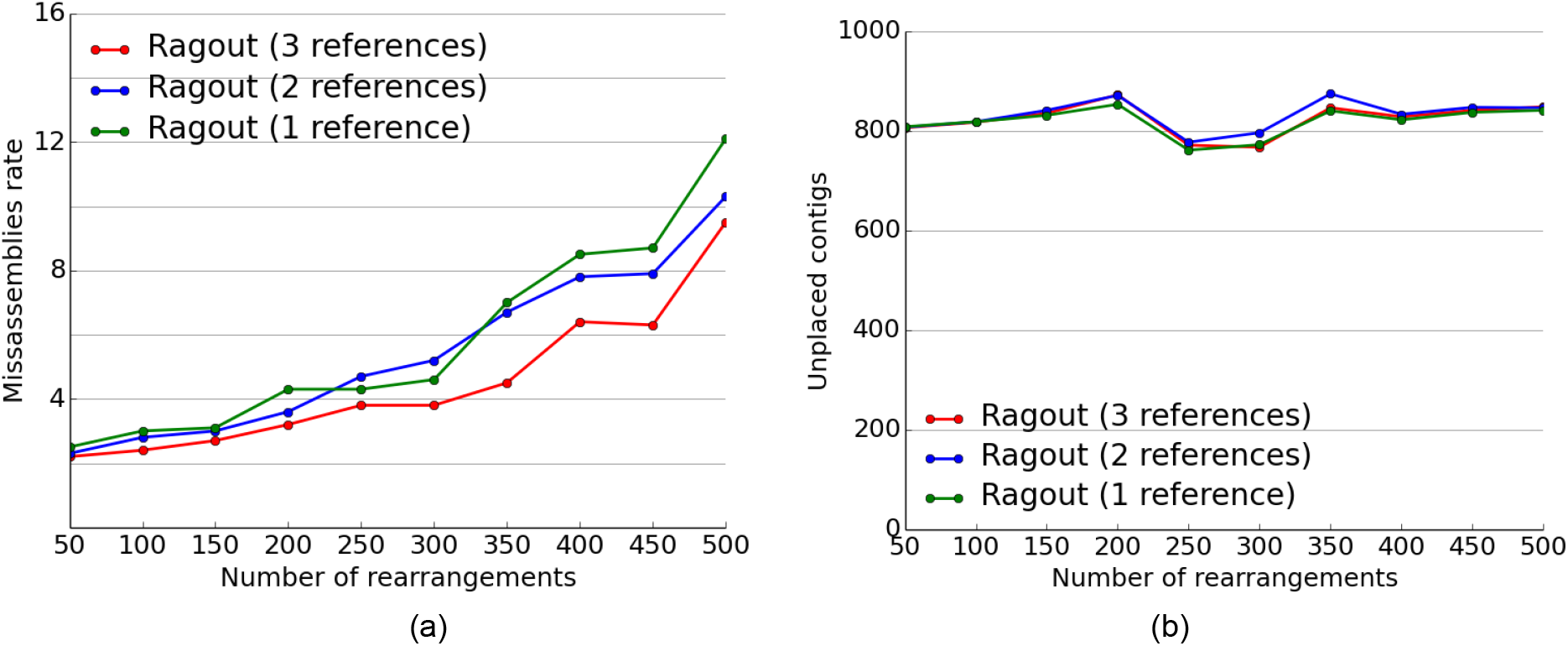
Misassembly rates (a) and number of unplaced contigs (b) for Ragout depending on the number of references used (with one closest, two closest and all three references). The total length of unplaced contigs does not exceed 4mb.

### S7 Theoretical estimates of the validated adjacencies rate

Given the genome of size 3 gbp and the reads of length 3,000 bp at 1x coverage, we estimated the probability of a region of length 500 bp (1,000 bp) is covered in full by at least one read as 56 % (51 %) through the direct modeling of read sampling. Thus, the probability of a correct and covered adjacency without a gap not validated by any reads could be copmuted as:

> *Prob(left flanking region of size 500 bp is covered) x*
>
> *Prob(right flanking region of size 500 bp is covered) x*
>
> *Prob(adjacency region of 1000bp is not covered)* ≈*15 %*

